# A framework to identify modifier genes in patients with Phelan-McDermid syndrome

**DOI:** 10.1101/117978

**Authors:** Anne-Claude Tabet, Thomas Rolland, Marie Ducloy, Jonathan Lévy, Julien Buratti, Alexandre Mathieu, Damien Haye, Laurence Perrin, Céline Dupont, Sandrine Passemard, Yline Capri, Alain Verloes, Séverine Drunat, Boris Keren, Cyril Mignot, Isabelle Marey, Aurélia Jacquette, Sandra Whalen, Eva Pipiras, Brigitte Benzacken, Sandra Chantot-Bastaraud, Alexandra Afenjar, Delphine Héron, Cédric Le Caignec, Claire Beneteau, Olivier Pichon, Bertrand Isidor, Albert David, Jean-Michel Dupont, Stephan Kemeny, Laetitia Gouas, Philippe Vago, Anne-Laure Mosca-Boidron, Laurence Faivre, Chantal Missirian, Nicole Philip, Damien Sanlaville, Patrick Edery, Véronique Satre, Charles Coutton, Françoise Devillard, Klaus Dieterich, Marie-Laure Vuillaume, Caroline Rooryck, Didier Lacombe, Lucile Pinson, Vincent Gatinois, Jacques Puechberty, Jean Chiesa, James Lespinasse, Christèle Dubourg, Chloé Quelin, Mélanie Fradin, Hubert Journel, Annick Toutain, Dominique Martin, Abdelamdjid Benmansour, Roberto Toro, Frédérique Amsellem, Richard Delorme, Thomas Bourgeron

## Abstract

Phelan-McDermid syndrome (PMS) is characterized by a variety of clinical symptoms with heterogeneous degrees of severity, including intellectual disability, speech impairment, and autism spectrum disorders (ASD). It results from a deletion of the 22q13 locus that in most cases includes the *SHANK3* gene. *SHANK3* is considered a major gene for PMS, but the factors modulating the severity of the syndrome remain largely unknown. In this study, we investigated 85 PMS patients with different 22q13 rearrangements (78 deletions, 7 duplications). We first explored their clinical features and provide evidence for frequent corpus callosum abnormalities. We then mapped candidate genomic regions at the 22q13 locus associated with risk of clinical features, and suggest a second locus associated with absence of speech. Finally, in some cases, we identified additional rearrangements at loci associated with ASD, potentially modulating the severity of the syndrome. We also report the first *SHANK3* deletion transmitted to five affected daughters by a mother without intellectual disability nor ASD, suggesting that some individuals could compensate for such mutations. In summary, we shed light on the genotype-phenotype relationship of PMS, a step towards the identification of compensatory mechanisms for a better prognosis and possibly treatments of patients with neurodevelopmental disorders.

## INTRODUCTION

Phelan-McDermid syndrome (PMS) is a severe medical condition characterized by hypotonia, global developmental delay, intellectual disability (ID), absent or delayed speech, minor dysmorphic features, and autism spectrum disorders (ASD).^1^ It results from a deletion of the 22q13 locus in the distal part of the long arm of chromosome 22. PMS occurs in 0.2-0.4% of individuals with neurodevelopmental disorders (NDD),^2^ but the prevalence remains difficult to ascertain due to important biases in the identification of the patients.

In most reported cases, the 22q13 deletion appeared *de novo^3^.* The size of the deleted genomic segment varies from hundreds of kilobases (kb) to more than nine megabases (Mb). The mechanisms resulting in the deletion are also highly variable, including simple deletions, unbalanced translocations, ring chromosomes or more complex chromosomal rearrangements.^2^ In the vast majority of the cases, the deletion includes *SHANK3,* a gene associated with several neuropsychiatric disorders including ID, ASD and schizophrenia.^4–6^ Rare interstitial deletions at 22q13 not encompassing *SHANK3* were also observed in patients with features of PMS, suggesting the implication of additional genes in that region or a positional effect influencing *SHANK3* expression.^7,8^

*SHANK3* codes for a scaffolding protein at the post-synaptic density of glutamatergic synapses, and is known to play a critical role in synaptic function by modulating the formation of dendrites.^9^ Mice lacking *Shank3* present alterations in the morphogenesis of the dendritic spines of hippocampal neurons and abnormal synaptic protein levels at the post-synaptic density of glutamatergic synapses.^10^ Studies using human neurons derived from induced pluripotent stem cells (iPSC) of patients with PMS revealed that abnormal networks are reversible by an overexpression of the SHANK3 protein or by treatment with insulin growth factor 1 (IGF1).^11^ *SHANK3* mutations can also cause a channelopathy due to a reduction of the hyperpolarization-activated cyclic nucleotide-gated channel proteins (HCN proteins).^12^ Finally, using iPSCs from patients with ASD and carrying *de novo SHANK3* loss-of-function mutations, Darville et al. identified lithium as a factor increasing *SHANK3* transcripts.^13^

Patients diagnosed with PMS display a large range of clinical features with various degrees of functional impact. Most patients are dependent on their caregivers. Associated medical or psychiatric comorbidities include seizures (grand mal seizures, focal seizures and absence seizures), renal abnormalities (mainly absent kidney, structural abnormalities of the kidney, hydronephrosis and kidney reflux), cardiac defects (tricuspid valve regurgitation, atrial septal defect, patent ductus arteriosus, and total anomalous pulmonary return), gastrointestinal disorders (intestinal/anal atresia, chronic constipation, gastroesophageal reflux), and ophthalmic features (most often strabismus).^14^ The molecular basis for this clinical heterogeneity is still largely unresolved. To date, at least 13 case series have been published gathering 584 affected patients with PMS.^14^ Only five studies (including 310 subjects) investigated the correlation of clinical features variability and the size of the 22q13 deletion,^15–19^ but the causality remains unclear. In addition, none of these studies explored the role of additional genetic variants (“multiple-hits”) in the diversity and severity of the symptoms reported in patients.^20,21^

In the present study, we collected clinical and genomic data from 85 patients carrying an unbalanced genomic rearrangement involving *SHANK3* (78 deletions and 7 duplications). We first describe the impact of the size of the 22q13 deletion on the clinical features. Then, we report the first identification of multiple-hits in patients with PMS, some affecting known ASD-risk loci that could influence the diversity and severity of the phenotype. Finally, we illustrate the clinical and genomic heterogeneity of individuals carrying 22q13 rearrangements in a family with five affected siblings who inherited a *SHANK3* deletion from a mother without ID nor ASD, providing a proof of principle that some individuals could compensate for such mutations.

## MATERIALS AND METHODS

### Population

Eighty-five patients (39 males, 44 females and two fetuses) carrying a genomic rearrangement at the 22q13 locus encompassing *SHANK3* were recruited through a French national network of cytogeneticists from 15 centers (ACHROPUCE network). For seven centers, the total number of microarray analyses performed for ID, congenital anomalies or ASD was available to establish a broad estimation of the prevalence of *SHANK3* deletions (Table S1). Detailed clinical information was collected for all subjects based on medical records, including perinatal events, growth parameters at birth and during early life of developmental, cognitive and functional development, congenital anomalies, dysmorphic features and main somatic comorbidities (Figure 1, Table S2, Supplementary Materials and Methods). Results from electroencephalograms (EEG) and brain magnetic resonance imaging (MRI) were also gathered for 21 and 35 individuals, respectively. We used the Diagnostic and Statistical Manual of Mental Disorder Fifth edition (DSM-5) criteria for ASD diagnostic, and standardized evaluations including the Autism Diagnostic Interview Revised (ADI-R)^22^ and the Autism Diagnostic Observation Schedule (ADOS-2).^23^ The cognitive level was also measured using Raven’s Progressive Matrices for nonverbal intelligence quotient (IQ) and the Peabody picture vocabulary test for verbal IQ. The local Institutional Review Boards (IRB) at each clinical center approved the study. For each patient, a parent or legal guardian consented to the study on his or her behalf.

**Figure 1.**
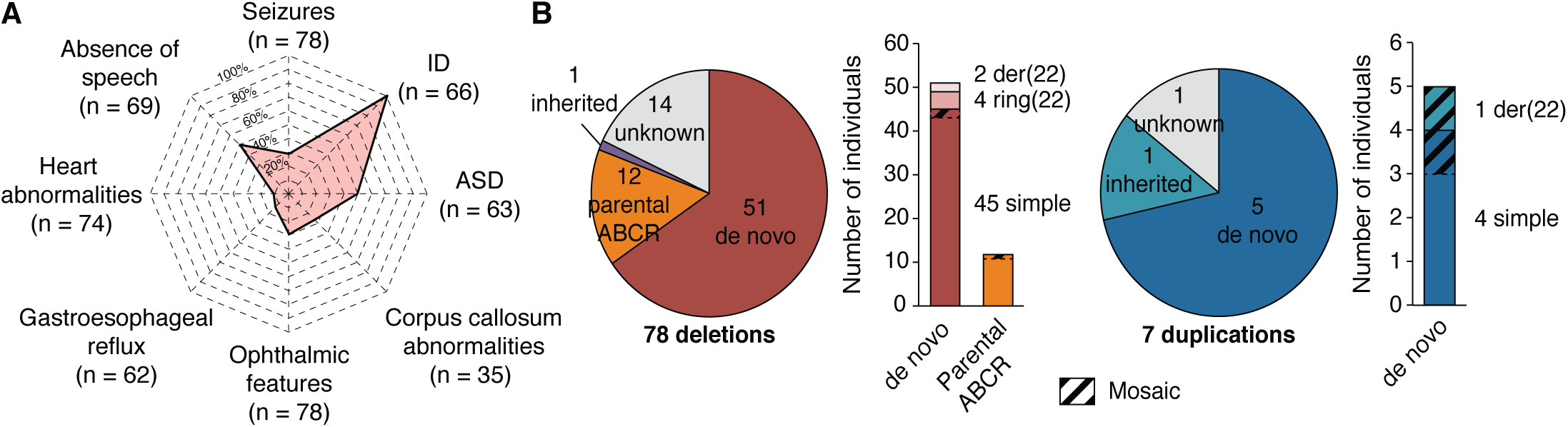
Clinical and genetic description of the cohort. (**A**) Spider plot of the clinical profile of the patients with PMS as a percentage of cases having each feature over those tested. (**B**) Mechanisms and inheritance of identified 22q13 CNVs in the cohort. ABCR: apparently balanced chromosomal rearrangement; der(22): derived from unbalanced reciprocal translocation; ring(22): caused by a ring chromosome 22.

### Identification of Genomic Rearrangements

Genomic data included information on the type of genomic rearrangement at the 22q13 locus, the methods of detection and confirmation, the molecular coordinates and the inheritance status (Table S3). We considered the size and breakpoints of the 22q13 rearrangement only for 74 patients studied using microarray technologies (Figure S1, Table S3).

Quality controls were performed locally in each genetic laboratory (Table S3). Genomic coordinates of each CNV are given according to the hg19 version of the human reference genome. All 22q13 rearrangements were validated in the corresponding clinical center using independent molecular technologies such as FISH, qPCR or MLPA. Inheritance of the 22q13 rearrangement was also determined in the corresponding laboratories.

We mapped all CNVs in the genomes of the patients, and focused on a list of 1,184 candidate genes for neuropsychiatry (NP-genes), which included Class I-III genes,^24^ TADA-65 genes,^25^ the genes from the SFARI database (release from the 9^th^ November 2015; https://gene.sfari.org), and the developmental brain disorder genes (Supplementary Tables 4 and 5).^26^ Four patients (P20, P22, P79 and P81) were not included in the analysis since their arrays did not pass the quality control for a whole-genome analysis. In 14 cases with unbalanced reciprocal translocations (Table S3), we considered the other chromosomal unbalanced segments as additional CNVs. The burden of CNVs in patients with PMS was compared to independent cohorts of patients with NDD (Figure S2).

### Computational and Statistical Analyses

Most analyses were performed on genetic and clinical data from 74 patients with array data (67 with a 22q13 deletion, 7 with a duplication), excluding 11 patients for whom only standard cytogenetic data were available (Table S3).

Ward’s hierarchical agglomerative clustering analysis was performed using JMP Pro 10.0.2 (http://www.jmp.com) on 40 individuals for whom data on ASD, speech, seizures and deletion size were available (Table S2).

For the mapping of the 22q13 locus, the prevalence of each feature was measured each 50 kb in sliding windows of 1.5 Mb. For each window, we calculated the percentage of individuals carrying a CNV starting in the corresponding interval and showing ASD, absence of speech, ophthalmic features, seizures, gastroesophageal reflux, heart abnormalities or corpus callosum abnormalities. Then, for each feature, we identified the interval defined by the minimal and maximal positions at which the prevalence was higher than the overall prevalence of the feature in the cohort. Finally, the association of the region with the feature was measured by a Fisher’s exact test for the enrichment odds ratio (OR) and p-value of individuals having the feature and carrying a CNV starting in the region compared to the rest of the 22q13 locus. To measure the probability to find such regions given the clinical and genetic sampling of the cohort, we developed a bootstrap-based measure of the statistical power. For each feature, we re-shuffled 1,000 times the feature presence/absence of the individuals, and reran our analysis of prevalence. We counted the number of times the enrichment OR was higher than the one observed with the original data, and used it as an empirical p-value (hereafter referred to as sampling p-value).

## RESULTS

### The 22q13 Rearrangements: Prevalence and Clinical Features

Based on the results of seven clinical centers gathering 18,115 patients with ID, ASD and/or congenital malformations, we could provide a broad estimation of 0.27% for the prevalence of 22q13 deletions (0.21 to 1.38% depending on the clinical center) (Table S1). Unlike most NDD for which males are significantly more affected than females, we observed a balanced sex ratio with 37 males and 39 females carrying a 22q13 deletion. Clinical features of patients included in this study were similar to those previously reported in larger datasets of patients (Figure 1, Supplementary Materials and Methods). However, for 35 patients, we analyzed the structural brain MRI data and 23 had macroscopic anomalies (65.7%), consistent with those already described in PMS (Supplementary Materials and Methods). Remarkably, 10 of these patients (28%) presented abnormalities of the corpus callosum ranging from thinness to agenesis (Figure S3) indicating that this abnormality is found in a relatively large subgroup of patients with PMS.

### The 22q13 Rearrangements: Inheritance and Sizes

Our cohort included 78 patients carrying a deletion at 22q13 overlapping *SHANK3* and seven patients with a duplication (Table S3). The deletions mostly derived from simple *de novo* deletions (57%), including two that were mosaic in the somatic cells of the patients (Figure 1). Other *de novo* chromosomal rearrangements included terminal deletions caused by a ring chromosome 22 in four cases, and deletions derived from unbalanced reciprocal translocations in two cases. Altogether, *de novo* rearrangements accounted for 51 cases (65.4%). In 12 cases, the deletions resulted from recombined parental balanced translocations or inversions. The size of the deleted segment was highly variable, ranging from 45.8 kb to 9.1 Mb (Figure S4).

The duplications occurred *de novo* in 5 cases. For P10, it was inherited from a healthy mother and for P61, inheritance was unknown. Two duplications were in mosaic (P29 and P50), including one case (P29) where the duplication resulted from an unbalanced translocation. The size of the duplicated segment ranged from 96 kb to 5.8 Mb.

### Exploratory Cluster Analysis of PMS Clinical Heterogeneity

To identify homogeneous groups of patients, we performed a hierarchical clustering analysis on 40 individuals for whom information concerning sex and presence or absence of ASD traits, speech and seizures were available. Five clusters summarized the variability (Figure 2). The main separation was based on the size of the deletion. Clusters C1 (six patients) and C2 (nine patients) were characterized by large deletions of an average of 5 and 7 Mb, respectively. These two clusters showed very similar percentages of patients with absence of speech (>80% in both clusters), but differed specifically for seizures with 0% and 67% of the patients in cluster C1 and C2, respectively. Patients with smaller deletions were divided in three clusters. Clusters C3 (eight patients) and C4 (ten patients) both had deletions of medium size (1.4-1.6Mb), but differed for sex and ASD (C3: 100% female and 0% ASD; C4: 100% males and 60% ASD). Finally, cluster C5 included seven patients with the smallest deletions and all with ASD and seizures. Therefore, the size of the 22q13 deletion might at least partly explain the presence and severity of the PMS symptoms, prompting a more detailed mapping of the genomic regions associated to specific symptoms in order to identify modifier genes.

**Figure 2.**
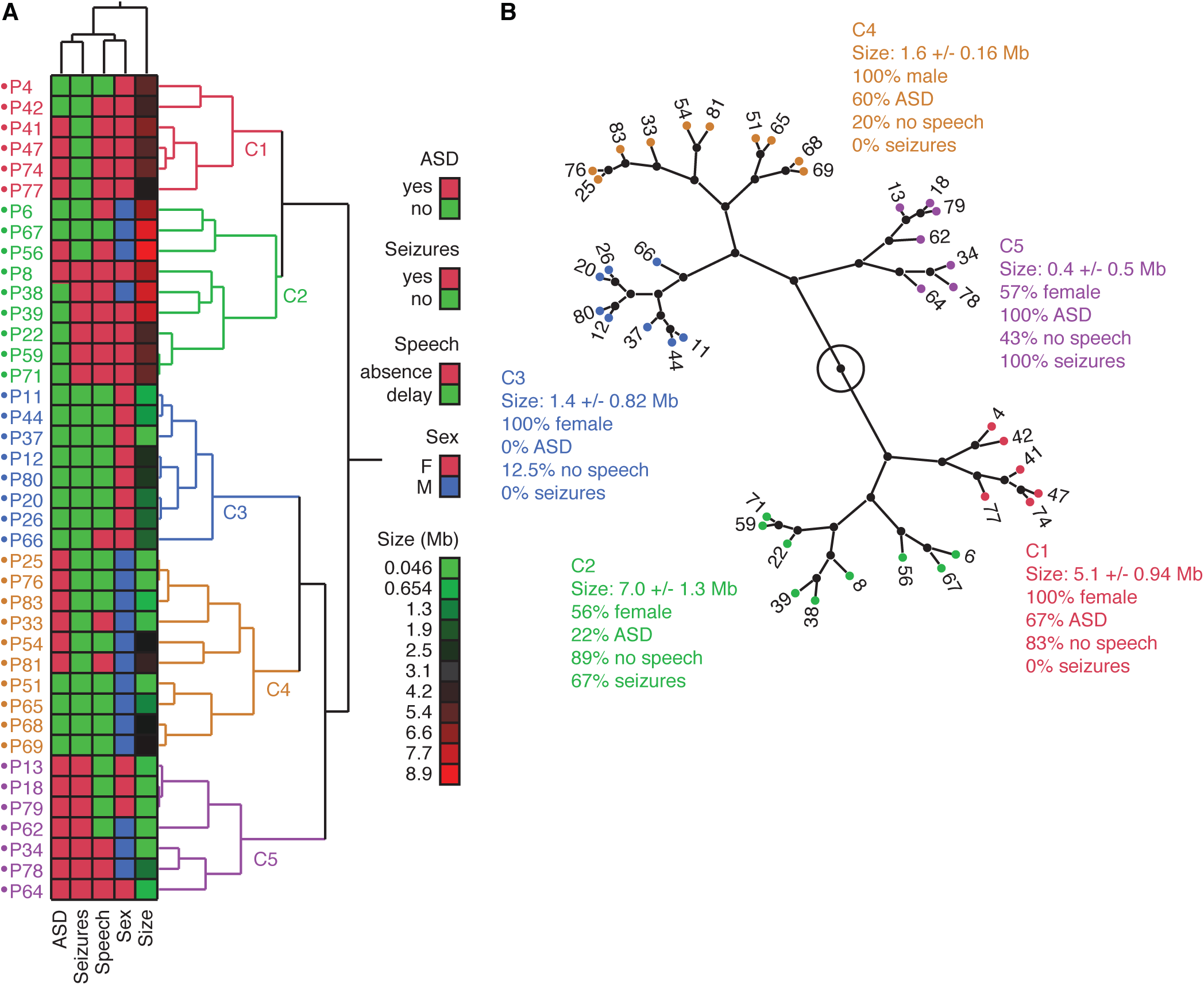
Multivariate analysis of the 22q13 deletion size, sex and clinical features of patients. (**A**) Hierarchical clustering based on multivariate analysis shows five clusters. (**B**) Tree view showing the five clusters of patients and the main genomic and clinical features of each cluster.

### Association between Clinical Features and Genomic Regions at 22q13

To map the genomic regions associated to PMS-related features, we measured the prevalence of ASD traits, absence of speech, ophthalmic features, seizures, gastroesophageal reflux, heart abnormalities or corpus callosum abnormalities in sliding windows of 1.5 Mb, each 50 kb in the 22q13 locus (Figure 3, Materials and Methods). Using this approach, we mapped candidate 22q13 regions conferring a high risk to display specific clinical features, which were statistically assessed for enrichment (odds ratio and p-value) and for statistical power to identify such a region given the sampling of the cohort (sampling p-value, Materials and Methods). We found regions with significantly higher risk for absence of speech, ophthalmic features and gastroesophageal reflux.

**Figure 3.**
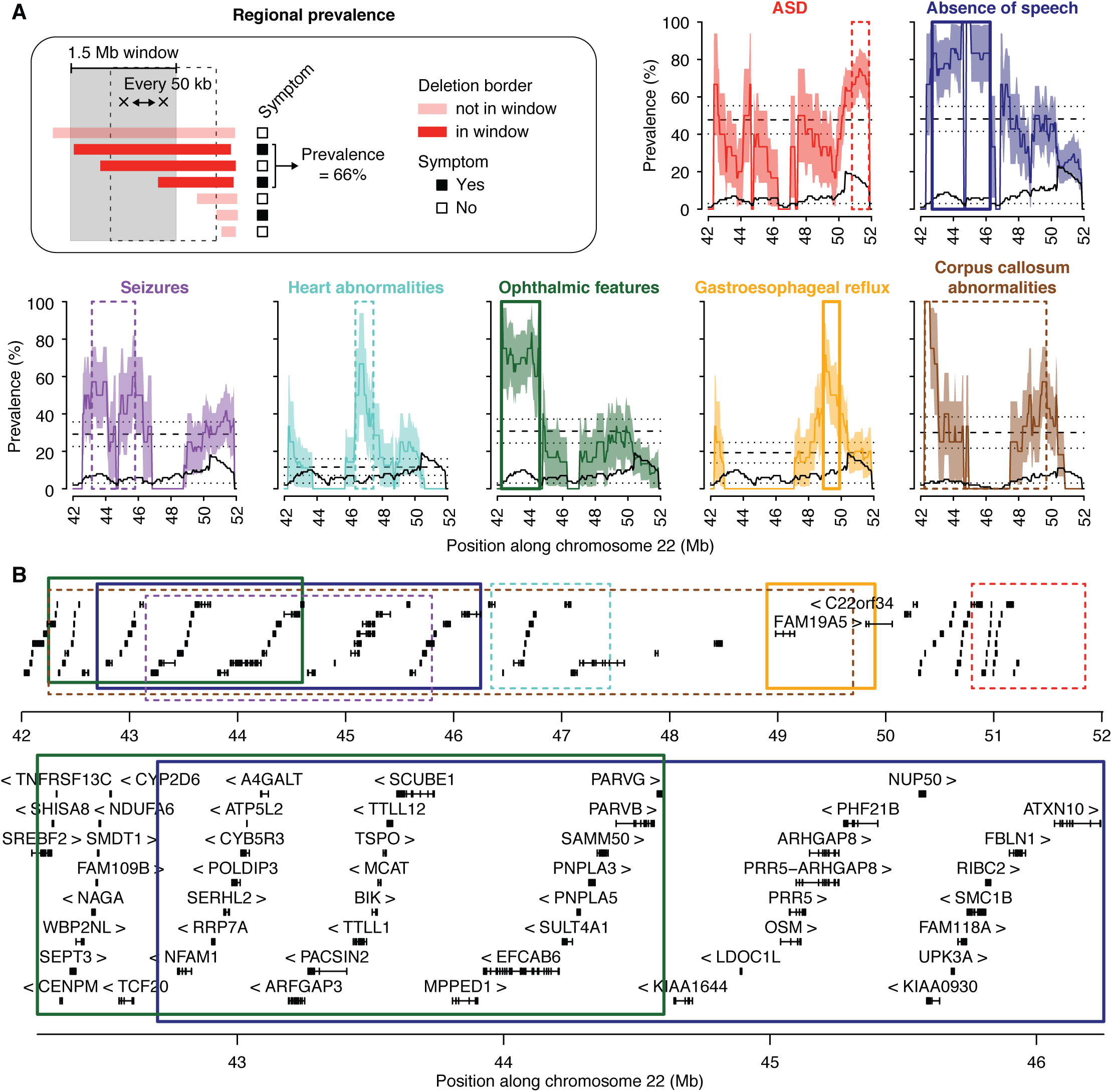
Mapping of genomic regions at 22q13 associated with high risk of presenting clinical features. (**A**) Prevalence is measured each 50 kb, within overlapping windows of 1.5 Mb (Materials and Methods). Dashed lines represent the global prevalence of each feature, measured as the fraction of patients with 22q13 deletions presenting the feature over all patients with 22q13 deletions. Dotted lines and colored areas represent standard errors of the proportion. Black solid lines show the numbers of informative patients for each window, and black dotted lines correspond to the minimum number of individuals required by window (n=3) to reduce interpretation biases. (**B**) The 22q13 locus is represented with the genes (block: exon, line: intron, arrow: strand) and the regions corresponding to a higher than global prevalence for each feature (top). Regions significantly associated with absence of speech and ophthalmic features are also shown in more details (bottom). Color code similar to (A).

The prevalence for absence of speech was 49±6% in the cohort, but increased to 80% for individuals who carry deletions including the genomic segment between position 42.7-46.25 Mb on chromosome 22 (enrichment OR= 6.7, P=0.006 and sampling P=0.039, Figure 3). Thirty-eight protein-coding genes are located within this region, including *PACSIN2, MPPED1, SULT4A1, ATXN10* and *PARVB* that are all expressed in the brain and represent compelling candidates for increasing the risk of absence of speech in individuals with PMS.

The overall prevalence of ophthalmic features was 30%, but increased to 70% when the deletions include the 42.25-44.6 Mb region (enrichment OR=10.3, P=0.002 and sampling P=0.041, Figure 3). Two members of the Parvin gene family, *PARVB* and *PARVG,* are located at the boundary of this region and code for actin binding proteins.

The prevalence of gastroesophageal reflux was 30% in the cohort, increased to 70% when deletions included the 48.9-49.9 Mb region (enrichment OR=12.3, P=0.009 and sampling P=0.041). This region contains the *FAM19A5* gene, encoding a small secreted protein mainly expressed in the brain and potentially acting as a modulator of immune response in nervous cells.^27^

For the remaining clinical features associated with PMS (ASD traits, heart abnormalities, seizures and corpus callosum abnormalities), no genomic regions were significantly associated with increased risk and supported by both enrichment and sampling statistical measures, suggesting that other loci or environmental factors could act as modifiers modulating the presence or severity of these features in PMS.

### Additional CNVs in Patients with PMS

In order to test whether other loci could modulate the severity of the clinical features, we systematically identified all the CNVs carried by the patients and affecting the exons of 1,184 candidate neuropsychiatric risk genes (hereafter referred as NP-genes, Table S4, Materials and Methods). Among the 63 patients carrying 22q13.3 deletions and tested using an array technology, 41 carried at least one CNV including at least one exon of a NP-gene (Figure 4, Figure S5, Table S5). Twenty-five patients had only one such CNV, nine had two, and seven had three or more. This burden of CNVs affecting NP-genes did not differ from the one measured in independent cohorts of patients with NDD and tested on similar array platforms (Figure S2). Interestingly, we found five patients who carried CNVs recurrently associated with ASD (Supplementary Materials and Methods). Patient P31 carried a 854 kb duplication at 16p11.2 inherited from a healthy mother, patient P53 carried a 494 kb deletion at 16p11.2 of unknown inheritance, patient P60 carried a 300 kb duplication at 15q11.2 of unknown inheritance, patient P67 carried a 2.7 Mb duplication at 16p12.3 of unknown inheritance, and patient P77 carried a 475 kb deletion at 15q11.2 inherited from a healthy father.

**Figure 4.**
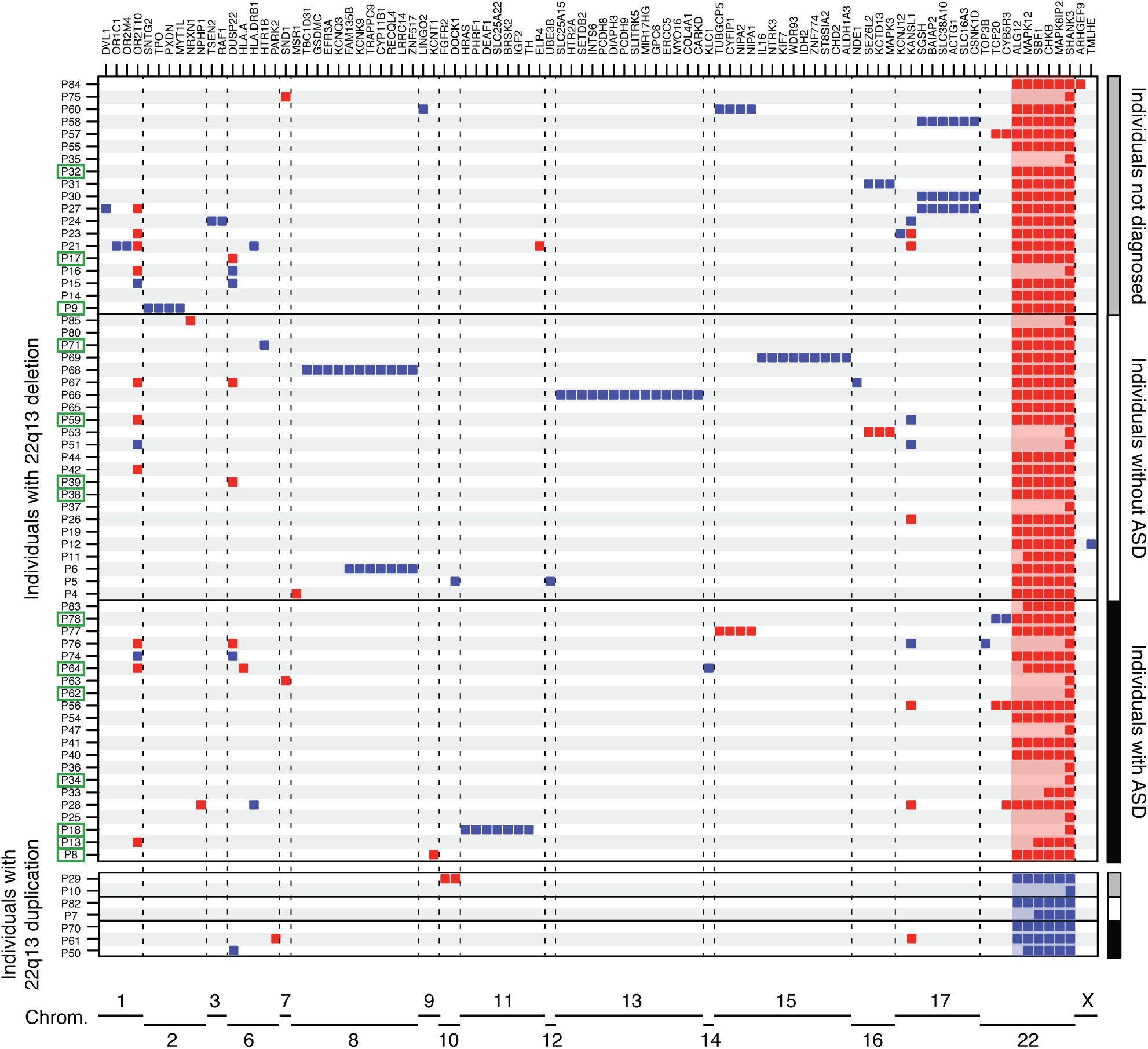
Additional CNVs identified in the cohort and including at least one NP-gene. Deletions (red) and duplications (blue) are represented for each patient and each NP-gene.

Out of the 14 patients carrying 22q13 deletions and having seizures, four were carrying an additional CNV covering a known risk gene for epilepsy: *KCNT1* (OMIM: 608167), *MYT1L* (OMIM: 613084), *DEAF1* (OMIM: 602635) and *SLC25A22* (OMIM: 609302) (Figure 4, Figure S5). For example, patient P8, who was severely affected and unable to speak, had a ring chromosome 22 resulting in a terminal 22q13 deletion of 7.1 Mb and an additional 87 kb deletion at 9q34 including *KCNT1* that codes for a sodium-activated potassium channel, largely expressed in the nervous system and implicated in autosomal dominant forms of epilepsy.

### A Multiplex Family with an Inherited *SHANK3* Deletion

Among the patients included in this study, we identified one female patient (subject III-D, Figure 5) who carried a maternally inherited *SHANK3* deletion of 67 kb, removing exons 1-8 of the isoform A. In addition, she also carried a 104 kb deletion of *NRXN1* at 2p16.3 (exons 3-5 of the Alpha 2 isoform and exons 3-4 of the Alpha 1 isoform). This proband was a member of a multiplex family that illustrates the heterogeneous clinical severity and the presence of multiple hits in the genome of the patients.

**Figure 5.**
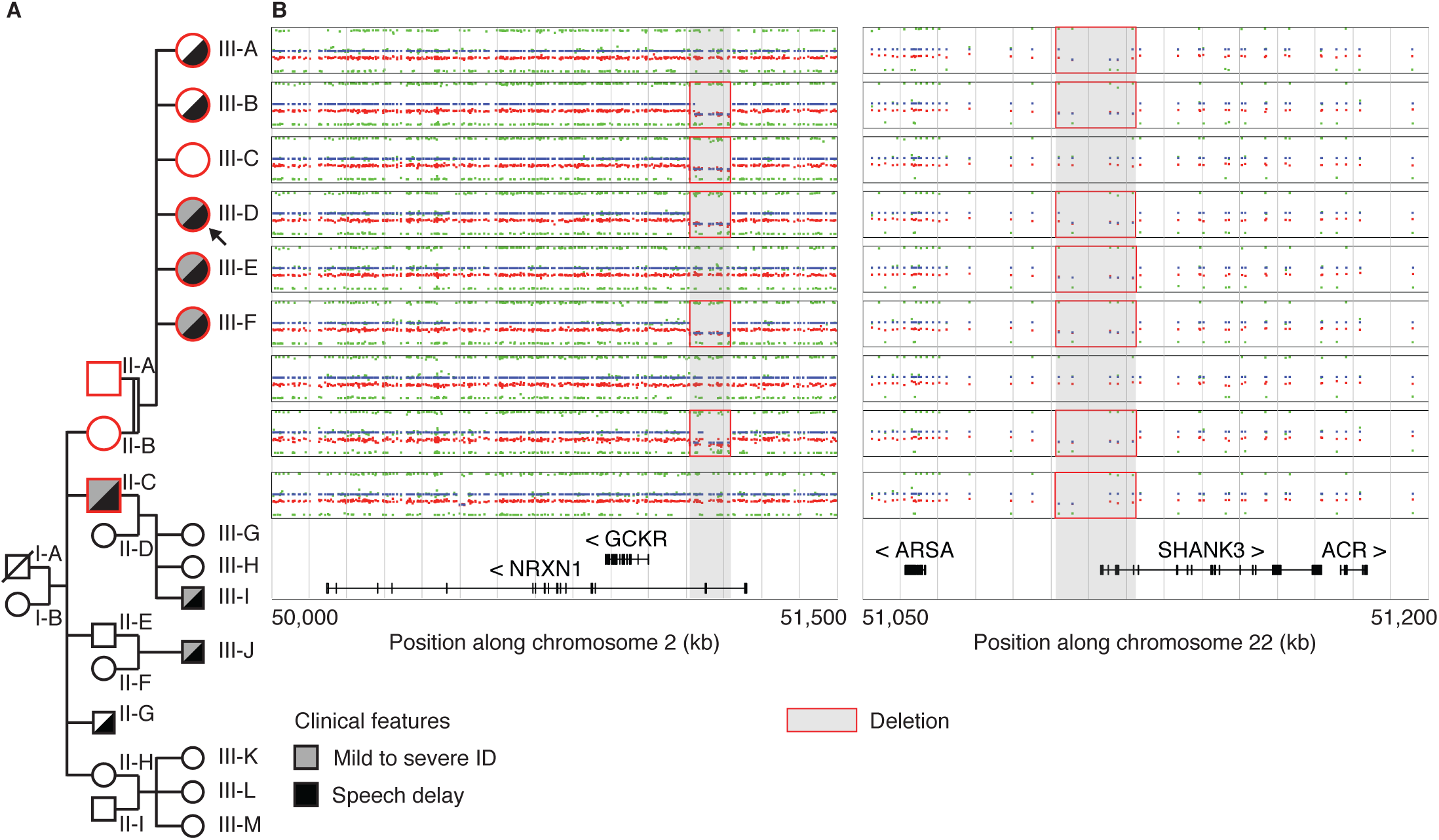
Multiplex family with inherited *SHANK3* and *NRXN1* deletions. (**A**) The nine studied individuals are displayed in the pedigree of the family. (**B**) CNVs detected by OmniExpress Illumina arrays and including the *NRXN1* and *SHANK3* genes (log2 scale).

The proband was a 10-year-old female, fourth child (among six) of consanguineous parents (2nd degree). Birth and neonatal parameters were in the normal range. Despite normal development during infancy, the parents reported paucity of social interactions, but without stereotyped body movements. The patient exhibited mildly delayed speech, with first words between three and four years old. At eight years old, she was diagnosed with ID, and assessed for cognitive impairment (full scale IQ=49, below the first percentile for her age). She did not meet criteria for ASD or any additional axis I psychiatric comorbidities. The patient presented with minor dysmorphic features including a curved forehead with a high implantation of her hair, a short and broad nose with flat nasal tip, long flat philtrum, a thick upper lip with a retrognathy. Concordant with her cognitive defects, she attended a special needs school and acquired basic writing and reading skills.

The father had no personal medical or psychiatric history and carried none of the two CNVs including *SHANK3* and *NRXN1.* In contrast, the mother (subject II-A, Figure 5) had mild learning difficulties related to a borderline IQ (non-verbal IQ=75), but she self-reported no history of speech delay. She carried the two deletions including *SHANK3* and *NRXN1.* The clinical assessment of the mother revealed that she had no significant autistic symptoms, no axis I psychiatric comorbidities and no significant medical history. She shared similar signs of dysmorphism with her daughter.

We also investigated the sisters of the proband (subject III-A, III-B, III-C, III-E and III-F; Figure 5, Supplementary Materials and Methods). Only one girl (III-C) did not carry the *SHANK3* deletion. She had a full scale IQ in the normal range (FS-IQ=92). She did not display any significant autistic symptoms (Social Responsiveness Scale total score < 66 percentile) or any signs of developmental delay (specifically no significant delay in speech development). The four other girls presented either moderate speech delay with first sentences at 4 years (III-A, III-B), or both speech disorder and mild to moderate ID (III-E, III-F). However, none of them displayed significant autistic symptoms, nor axis I psychiatric comorbidities and significant medical history. Finally, we investigated one of the maternal uncles (subject II-C, Figure 5), who also carried the *SHANK3* deletion and presented mild ID associated with a severe language impairment. He currently has a job dedicated to persons with mental health problems.

Genetic testing showed that the five affected daughters and the maternal uncle carried the 22q13 deletion removing the first part of isoform A of *SHANK3.* In addition, subjects III-B, III-C, III-D, III-F and II-C also had the 2p16.3 deletion overlapping *NRXN1* (Figure 5). All affected subjects carried the partial *SHANK3* deletion suggesting its pathogenicity in the syndrome. The 2p16.3 deletion coexists in three patients with the 22q13 deletion, two with ID and speech delay (III-D and III-F) and one with only speech delay (III-B). This CNV could act as a modifier factor contributing at least in part to the clinical variability of the syndrome in this family. The unaffected girl (III-C) carried only the *NRXN1* deletion, suggesting that, alone, this CNV should not be considered as a risk factor.

## DISCUSSION

### Prevalence of PMS, ASD, Seizures and Brain Structural Abnormalities

Our broad estimation of the prevalence of patients with PMS in patients with NDD is 0.27% (0.21-1.38%), which is similar to the ones reported previously (0.2-0.4%).^2^ We also observed a balanced sex ratio as previously reported.^1^ Regarding the clinical features (Table S2, Supplementary Materials and Methods), we confirmed previous reports: in most cases, patients with PMS received a diagnosis of NDD, ID and/or speech impairment.

In our cohort, 50% of the patients were reported with autistic traits. In the literature, the frequency of patients with PMS diagnosed with ASD varied significantly from one study to another mostly because of inconsistent methods of evaluation. In a report of eight patients, Philippe et al. (2008) did not diagnose ASD in any of the patient.^28^ In contrast, Phelan et al. (2012) diagnosed ASD in 17 out of 18 patients with PMS.^1^ Soorya et al. (2013) showed that 27 out of 32 patients with PMS (84%) have a diagnosis of ASD when standardized methods are used.^16^ Based on standardized interview with parents/caretakers, a recent study estimated that approximately 50% of PMS patients met criteria for ASD (21 patients out of 40).^29^ Similarly, the prevalence for seizures seems highly heterogeneous among studies since it ranges from 17 to 70% in multiple case series.^30^ In Kolevson et al. (2014), 25% of the patients from 13 independent studies (121/482) had seizures,^14^ a frequency very similar to this cohort (24%;19/78) (Supplementary Materials and Methods).

In the literature, brain structural abnormalities were reported in six studies, with a mean rate of 25% (58/233) of cases.^14^ Hypoplasia of the cerebellar vermis,^31,32^ thin corpus callosum,^16,28,31,33^ and abnormalities of white matter were previously reported. In our cohort, brain MRI data was available for 35 patients with 22q13 deletions and 23 had structural anomalies (65.7%), consistent with those already described in PMS (Supplementary Materials and Methods).

### Impact of the 22q13.3 Deletion Size on the Absence of Speech

We observed that ASD was associated with smaller deleted segments, and could confirm that absence of speech was associated with large deletions.^15,18^ Similarly, Sarasua et al. (2011) reported that patients with an absence of speech had deletion breakpoints at 43.9 Mb (22q13.2) or more proximal.^15^ In our study, patients with breakpoint proximal to 46 Mb were more at risk for absence of speech than patients with smaller distal deletions (Figures 2 and 3). Within the 42.7-46.25 Mb interval, several genes are expressed in the brain and are compelling candidate for increasing the risk of absence of speech. *PACSIN2* (protein kinase C and casein kinase substrate in neurons 2) is a member of the pacsin-syndapin-FAP52 gene family and is upregulated upon neuronal differentiation. It has an essential role in the organization of clathrin-mediated membrane endocytosis in neurons.^34^ The function of *MPPED1* remains uncharacterized, but it codes for a metallophosphoesterase domain-containing protein highly expressed in the human fetal brain. *SULT4A1* codes for a protein of the sulfotransferase family involved in the metabolism of endogenous factors such as hormones, steroids, and monoamine neurotransmitters, as well as drugs and xenobiotics. *SULT4A1* is exclusively expressed in neural tissues, is highly conserved, and has been identified in all vertebrate studied so far. Interestingly, zebrafish carrying homozygous *SULT4A1* mutations exhibit excessively sedentary behavior during the day,^35^ suggesting a role of this gene in regulating behavior. The function of *ATXN10* remains largely unknown, but expanded ATTCT pentanucleotide repeats in intron 9 of the gene cause a rare form of spinocerebellar ataxia (SCA10) characterized by cerebellar ataxia and epilepsy.^36^

Interestingly, the region associated with absence of speech did not include *WNT7B,* a member of the WNT signaling molecule involved in the formation of the central nervous system vascular endothelium^37^ as well as in dendrite development.^38^ In contrast, this region includes the minimal interval of interstitial 22q13 deletions (not involving *SHANK3)* causing clinical features common to PMS^7^ and including *SULT4A1* and *PARVB. PARVB* is not highly expressed in the brain, but it codes for an actin-binding protein that interacts with ARHGEF6, a protein coded by an X-linked gene mutated in patients with ID.^39^

### Multiple Hits in Patients with PMS

To our knowledge, this is the first report investigating the presence of multiple-hits in patients with PMS. Among the 63 patients with array results, 41 carried at least one additional CNV encompassing exonic sequences of one NP-gene. It is important to consider that a duplication of a gene might not be causative and therefore CNVs affecting NP-genes might not always be deleterious. Nevertheless, we found 16p11.2 CNVs (one deletion and one duplication) in two independent patients (P32 and P59), which are known to increase risk of NDD and are rare in the general population (0.03%).^2^ The finding of two cases with such CNVs is intriguing and raises question of the genetic architecture of PMS, and by extension on the cumulative genetic risk factors in NDD.^40^ In addition, we observed abnormal gene dosage of risk-genes for epilepsy such as *KCNT1, MYT1L, DEAF1* and *SLC25A22* in patients whom had lifetime history of seizures. However, at that stage, we cannot conclude on the causative effect of such CNVs since we also observed patients with CNVs affecting risk-genes for epilepsy (such as *KANSL1* or *NRXN1),* and a lifetime absence of epilepsy. In summary, multiple-hits exist in patients with PMS, but their impact on the severity of the symptoms remains to be determined in larger cohorts of well-phenotyped patients with more extensive genetic profiles using whole exome/genome sequencing.

### Conclusions

Our study confirms previous findings regarding the impact of the 22q13 deletion on several clinical features such as absence of speech and ASD. We also identified one mother without ID nor ASD carrying a *SHANK3* deletion, providing the first proof of principle that some individuals could be resilient for such mutations. Larger cohorts of individuals carrying 22q13 deletions (including interstitial deletions not affecting *SHANK3)* with in-depth phenotyping and whole genome sequencing data should allow us to identify modifier genes. Such genes and pathways would participate to our understanding of the etiology of PMS and could represent new relevant drug targets.

## ACKNOWLEDGEMENTS

We are grateful to the families for participating in this study. We thank Isabelle Cloëz-Tayarani, Claire Leblond, Ksenia Bagrintseva and Gaël Millot for helpful discussions and careful reading of the manuscript.

The authors declare no competing interests.

